# A high-throughput radioactivity-based assay for screening SARS-CoV-2 nsp10-nsp16 complex

**DOI:** 10.1101/2021.02.03.429625

**Authors:** Aliakbar Khalili Yazdi, Fengling Li, Kanchan Devkota, Sumera Perveen, Pegah Ghiabi, Taraneh Hajian, Albina Bolotokova, Masoud Vedadi

## Abstract

Frequent outbreaks of novel coronaviruses (CoVs), highlighted by the current SARS-CoV-2 pandemic, necessitate the development of therapeutics that could be easily and effectively administered world-wide. The conserved mRNA-capping process enables CoVs to evade their host immune system and is a target for antiviral development. Nonstructural protein (nsp) 16 in complex with nsp10 catalyzes the final step of coronaviral mRNA-capping through its 2’-O-methylation activity. Like other methyltransferases, SARS-CoV-2 nsp10-nsp16 complex is druggable. However, the availability of an optimized assay for high-throughput screening (HTS) is an unmet need. Here, we report the development of a radioactivity-based assay for methyltransferase activity of nsp10-nsp16 complex in a 384-well format, and kinetic characterization, and optimization of the assay for HTS (Z′-factor: 0.83). Considering the high conservation of nsp16 across known CoV species, the potential inhibitors targeting SARS-CoV-2 nsp10-nsp16 complex may also be effective against other emerging pathogenic CoVs.

## 1. Introduction

The recent outbreak of SARS-CoV-2 (COVID-19) has posed a remarkable threat to global public health and economy ^1^. CoVs are major viral pathogens of humans and animals, including bats, livestock, and numerous wild animals ^2, 3^. Among recognized CoV species ^1^, only seven are documented to infect humans. SARS-CoV, MERS-CoV, and SARS-CoV-2 cause severe symptoms in humans, while 229E, HKU1, OC43, and NL63 are associated with mild symptoms ^2^, ^4^. CoVs belong to the family of Coronaviridae. Their genome is a single-stranded RNA, which encodes 16 functional <u>n</u>on-<u>s</u>tructural <u>p</u>roteins (nsp1 to nsp16) and four main structural and accessory proteins ^5^. Most non-structural proteins are involved in synthesis and processing of CoV RNAs ^6^.

Several endogenous 5′-mRNA-cap structures are identified in mature eukaryotic cells that help recruiting various cellular factors to support efficient translation and improve mRNA stability ^7^, ^8^. Additionally, mRNA-caps provide a molecular platform to differentiate between self and foreign mRNAs ^9^, which could result in initiation of host immune responses when viral mRNAs are detected. In return, viruses have evolved an array of systems to combat these restrictions ^10^, ^11^. Viral mRNAs that maintain both the 7-methylguanosine cap (N7-meGpppN; called Cap-0) and 2′-O-methylation of the first nucleotide (N7-meGpppN_2′Ome_; Cap-1) could stay viable like the host mRNAs ^12, 13^. Particularly, CoVs generate mRNAs with a type-1 cap structure to avoid identification and activation of host defence mechanisms ^13-16^. Coronaviral mRNA-capping starts with the removal of 5′-γ-phosphate from nascent viral RNA by nsp13. A Guanosine monophosphate is then attached to the 5′-diphosphate by an RNA-guanylyltransferase to form GpppN-RNA. Subsequently, nsp14 methylates N^7^ of guanosine, giving rise to a Cap-0. Ultimately, Cap-0 is transformed into a doubly methylated (Cap-1) structure by nsp16 ^9^, ^17^.

The importance of conserved nsp16 for function and survival of CoVs has been documented *in vivo* and *in vitro* ^13^, ^16, 18, 19^. Nsp16 is a member of the 2′-O methyltransferase (MTase) family, catalyzing the transfer of a methyl group from S-adenosyl methionine (SAM) to RNA substrates ^20^. MTases are generally druggable with several highly selective and cell-active inhibitors of human MTases available ^21^. The *in vitro* 2′-O-MTase activity of nsp16 has been reported for Feline-CoV, MERS-CoV, and SARS-CoV ^22-25^. However, nsp16 was significantly active only when in complex with nsp10 ^23^. Nsp16 alone is unstable, and nsp10-nsp16 complex formation is essential for its binding to SAM and RNA substrate ^24^. The crystal structure of nsp16 has only been determined in complex with nsp10 ^26^. Several structures of nsp10-nsp16 from various CoV species are available as apo, and in complex with RNA substrate, SAM, or SAM analogues, which vastly enables the structure-based hit optimization ^24, 26-30^. Nsp10-nsp16 complex selectively binds and methylates longer CoV mRNAs and synthetic small RNAs with Cap-0 ^23^. Moreover, SARS-CoV nsp10-nsp16 methylates N7-meGpppA-RNA, but not N7-meGpppG-RNA, which provides some selectivity over the host mRNAs ^24^.

These studies indicate that the conserved nsp10-nsp16 complex is essential for CoVs ability to mimic the host mRNAs needed for viral replication ^12,13, 23^. Therefore, inhibition of nsp10-nsp16 complex activity could potentially hinder the pathogenesis of CoVs through eliciting a host immune response ^13, 15, 16^. However, availability of an optimized assay suitable for high-throughput screening (HTS) is an unmet need. Here, we report development and optimization of a scintillation proximity assay (SPA) for testing RNA MTase activity of nsp10-nsp16 complex, kinetic characterization, and high-throughput screening.

## 2. Materials and Methods

### 2.1. Reagents

Biotinylated RNA substrate (5′ N7-meGpppACCCCC-biotin) was synthesised by bioSYNTHESIS (Levisville, Texas, USA). 384- and 96-well Streptavidin PLUS High-Capacity FlashPlates, ^3^H-SAM, and ^3^H-biotin were from PerkinElmer (Massachusetts, USA). SAM, sinefungin, and SAH were from Sigma, Missouri, USA. SAM2^®^ Biotin-Capture Membrane was obtained from Promega, Wisconsin, USA. All reaction buffers contained 0.4 U/μL RNaseOUT™ ribonuclease inhibitor (Invitrogen, Massachusetts, USA).

### 2.2. Protein Expression and Purification

Expression and purification of SARS-CoV-2 nsp10-nsp16 is recently described.^31^ Briefly, nsp16 (S_1_-N_298_) and nsp10 (A_1_-Q_139_) were separately expressed in *Escherichia coli* BL21(DE3) RIL and purified to near homogeneity. The nsp10-nsp16 complex was prepared using the purified proteins in an 8 (nsp10) to 1 (nsp16) molar ratio, dialyzed in storage buffer containing 50 mM Tris-HCl (pH 8.0), 200 mM NaCl, 0.5 mM TCEP, and 5% glycerol and flash frozen.

### 2.3. Optimization of nsp10-nsp16 Assay Conditions

The initial MTase reactions were performed in a buffer similar to the reported condition for SARS-CoV nsp10-nsp16 complex ^23^ with some modifications. Accordingly, 10 μL mixtures containing 50 mM Tris (pH 8.0), 1 mM MgCl_2_, 5 mM DTT, 2 µM RNA substrate, and 250 nM nsp10-nsp16 complex were prepared. The reactions were started by addition of 4 µM SAM (16% ^3^H-SAM). Reactions proceeded for 1 hour, and then quenched by adding 10 μL of 7.5 M Guanidine hydrochloride followed by 60 μL of 20 mM Tris-HCl (pH 8.0). The reaction products were transferred into Streptavidin-coated FlashPlates for scintillation counting using a TopCount instrument (PerkinElmer, Massachusetts, USA). Reaction mixtures were prepared in triplicate. For determining the optimum buffer pH, 50 mM Tris-HCl was used for generating the pH profile ranging from 6.5 to 9.0. The effect of various reagents such as salts, detergents, reducing agents, BSA, EDTA, and DMSO was investigated through titration of each reagent in assay buffer at p 7.5 and measuring their relative activity compared to the control (i.e., reactions without additive) using the SPA-based assay. The following buffer was chosen as the optimal reaction condition: 50 mM Tris-HCl, 100 mM KCl, 1.5 mM MgCl_2_, 0.01% Triton-X-100, 0.01% BSA, and 5 mM DTT. All subsequent experiments were performed using this buffer condition. All reactions were performed at room temperature (23 °C).

### 2.4. Determination of Kinetic Parameters for nsp10-nsp16 Complex

For determining the kinetic parameters, reactions were carried out using the optimized buffer condition in triplicate in standard 96-well polypropylene plates. For each experiment, the concentration for one substrate (i.e., SAM or RNA) was varied, while the concentration of the second substrate was kept at near saturation (>3.5x *K*_*m*_). After starting the reaction by adding ^3^H-SAM, samples were taken at different time points, which were quenched by adding an equal volume of 7.5 M Guanidine hydrochloride. The level of ^3^H-methylated-RNA in each reaction was quantified by measuring the radioactivity level (CPM; counts per minute) employing both a high-capacity biotin-capture membrane-based approach and an SPA-based method. For the membrane-based method, 12 μL of the quenched mixture was spotted on streptavidin-coated SAM2^®^ Biotin-capture membranes. The membranes were washed twice with 2M NaCl (2 min each), rinsed with water three times (2 min each), and were allowed to fully air dry. Subsequently, the spotted squares were cut from membrane and placed separately into scintillation vials containing scintillant solution, MicroScint™-O (PerkinElmer, Massachusetts, USA). Finally, the ^3^H-methyl-RNA product was quantified using a Tri-Carb scintillation counter, PerkinElmer. In the SPA-based approach, the quenched mixtures were transferred into 96-well FlashPlates after adding 150 μL of 20 mM Tris-HCl pH8.0 in each well. To avoid saturation of beads in these plates with excess RNA substrate, only 6 μL of the quenched mixture was added to each well. ^3^H-Biotin at different concentrations were used as control. After overnight incubation, the level of ^3^H-methylated-RNA was measured by scintillation counting. Afterwards, the initial velocities were obtained from the linear portions of the reaction progression curves. The kinetic parameters were calculated using Michaelis-Menten equation by GraphPad Software. For clarity, when we report the activity of the protein complex as nmoles/min/mg, the “mg” refers to “mg of nsp16”. Since the complex is 1 (nsp16): 8 (nsp10), the molarity of nsp16 and the nsp10-nsp16 complex are the same.

### 2.5. Z′-Factor Determination

The quality and robustness of the nsp10-nsp16 assay was verified by the standard Z′-factor determination ^32^. Optimized reaction mixture containing 125 nM nsp10-nsp16 complex, and 0.8 μM RNA were prepared in the presence or absence of 200 μM sinefungin in 384-well format using an Agilent Bravo automated liquid-handling robot. Final DMSO concentration was 1%. The reactions were started by addition of 1.7 μM SAM (30% ^3^H-SAM) and were incubated for 30 minutes at 23 °C. After measuring signal by SPA-based method, the Z′-factor was calculated as previously described ^32^.

### 2.6. Screening a Collection of Chemical Probes

The library of 76 epigenetics chemical probes was from Structural Genomics Consortium (SGC; https://www.thesgc.org/chemical-probes/epigenetics). The compounds were screened at 50 μM with a final DMSO concentration of 1% in 125 nM nsp10-nsp16 complex, 0.8 μM RNA, and 1.7 μM SAM (30% ^3^H-SAM). Reactions containing 50 μM SAH and 1% DMSO were used as positive and negative controls, respectively. After 30 min incubation, reactions were quenched, transferred into SPA plates, and the incorporated radioactivity was quantitated as described above.

### 2.7. Sequence Analysis

Nsp16 protein sequences were taken from the CoV *ORF1ab* sequences accessible through UniProt database. These sequences consisted of 229E (<u>P0C6X1</u>), HKU1 (<u>P0C6X3</u>), NL63 (<u>P0CX5</u>), OC43 (<u>P0C6X6</u>), MERS-CoV (<u>K9N7C7</u>), SARS-CoV (<u>P0C6X7</u>), and SARS-CoV-2 (<u>P0DTD1</u>). The nsp16 sequences were aligned using Clustal Omega ^33^, and sequence similarities and secondary structure features were rendered by ESPript Version 3.0 ^34^. The sequence conservation among these sequences was mapped onto the crystal structure of nsp10-nsp16 from SARS-CoV-2 (PDB: <u>6WKS</u>) using Chimera Version 1.14 ^35^.

## 3. Results

### 3.1. Assay Development and Optimization

*In vitro* activity of SARS-CoV-2 nsp10-nsp16 complex was tested by monitoring the transfer of ^3^H-SAM to the biotinylated N7-meGpppACCCCC RNA substrate. The methylated RNA product was captured using SPA plates followed by recording the changes in CPM. Initial tests at 250 nM of nsp10-nsp16 complex, 2 μM RNA substrate, and 5 μM SAM indicated the protein complex is active with significant signal-to-noise ratio. The assay conditions were further optimized with respect to the pH of the buffer and the presence of several commonly used additives (**Fig. 1**). The complex was most active at pH 7.5 (**Fig. 1A**). Using this optimal pH, the effects of other buffer components were investigated. Although NaCl over a wide range of concentrations (10-100 mM) reduced the enzyme activity by about 30%, KCl had little effect on nsp10-nsp16 complex activity up to 100 mM, and MgCl_2_ slightly increased the signal (**Figs. 1B-D**). However, presence of Triton X-100 as low as 0.002% increased the signal by more than 20% (**Fig. 1E**). Tween-20 had a similar effect (**Suppl. Fig. 1A**). The reducing agents, TCEP and DTT, had no significant effect on enzyme activity (**Figs. 1G-H**). The presence of BSA at concentrations higher than 0.02% reduced the signal readout (**Fig. 1F**). EDTA at concentrations as low as 50 µM considerably reduced the activity (**Suppl. Fig. 1B**). Based on these observations, 50 mM Tris-HCl, 100 mM KCl, 1.5 mM MgCl_2_, 0.01% Triton X-100, 0.01% BSA, and 5 mM DTT was selected as the optimized buffer condition for SARS-CoV-2 nsp10-np16 complex MTase activity assays. Overall, the assay optimization led to 70% increase in assay signal over the starting assay conditions (**Suppl. Fig. 1C)**. The nsp10-nsp16 complex activity under the optimized conditions was not affected by DMSO up to 5% (**Fig. 1I**).

**Figure 1.**
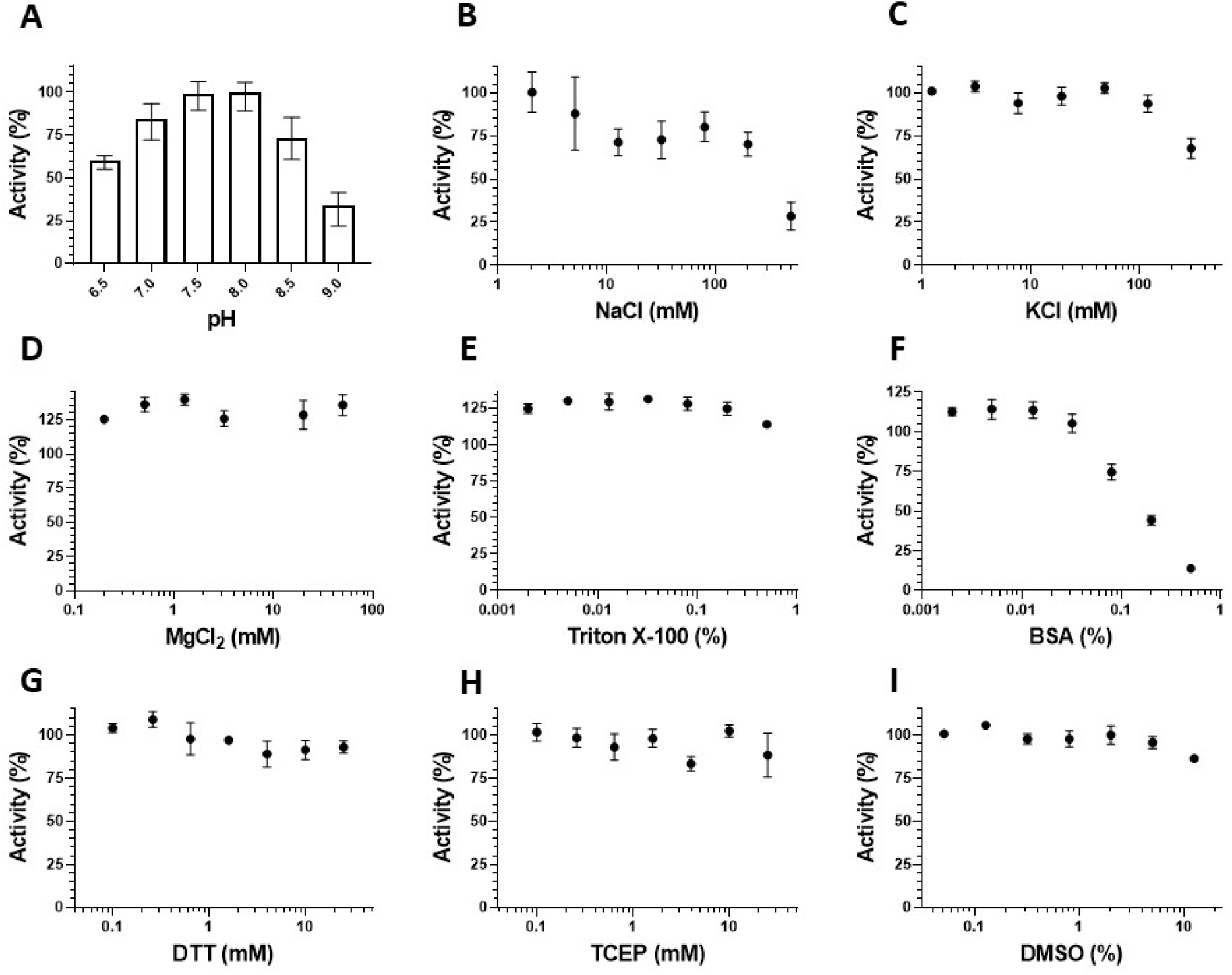
Assay optimization for the SARS-CoV-2 nsp10-nsp16 complex activity. **(A)** The methyltransferase activity of nsp10-nsp16 complex was tested at various pH. Tris-HCl was used to generate the pH gradient from 6.5 to 9.0. Using the optimal pH (7.5), the effects of **(B)** NaCl, **(C)** KCl, **(D)** MgCl_2_, **(E)** Triton X-100, **(F)** BSA, **(G)** DTT, **(H)** TCEP, and **(I)** DMSO were evaluated. Experiments were performed in triplicate. Values are mean ± standard deviations.

### 3.2. Kinetic Characterization

The kinetic parameters for nsp10-nsp16 complex were determined using the optimized conditions. Initial assessment of the MTase activity at various concentrations of nsp10-nsp16 complex indicated reaction linearity up to around 250 nM of the protein complex (**Supp. Fig. 1D**). At 250 nM of nsp10-nsp16 complex, using the membrane-based approach, apparent *K*_*m*_ values of 1.7 ± 0.3 μM and 1.6 ± 0.4 μM were determined for SAM and RNA, respectively, with apparent *k*_*cat*_ of 15.9 ± 1.2 h^-1^ (**Figs. 2A-B**). For determining the *K*_*m*_ of SAM, the concentration of RNA was kept at 5.6 μM, whereas when assessing the *K*_*m*_ of the RNA substrate, SAM concentration was at 6.0 μM. To investigate if lowering the concentration of nsp10-nsp16 complex is possible without nsp16 inactivation due to complex dissociation, the kinetic parameters for SAM and RNA were also determined at 125 nM of nsp10-nsp16 complex. Using a SPA, the linear initial velocities were used to calculate the kinetic parameters (**Suppl. Figs. 2A-B)**. The apparent *K*_*m*_ of 2.0 ± 0.2 μM and 1.0 ± 0.1 μM for SAM and RNA substrate respectively were determined (**Figs. 2C-D**). The apparent *k*_*cat*_ value was 26.9 ± 0.3 h^-1^. In this round of experiments, the concentration of the second substrates, SAM and RNA, were kept at 8.0 μM and 5.0 μM, respectively. These data indicated that nsp10 and nsp16 stay in complex at lower concentration and the integrity of the complex was not affected by further dilution of the protein complex. Therefore, all further assays were performed at 125 nM of nsp10-nsp16 complex. The N7-unmethylated biotinylated RNA substrate, which was used as a control, showed almost no activity under similar assay conditions (**Suppl. Fig. 1E**).

**Figure 2.**
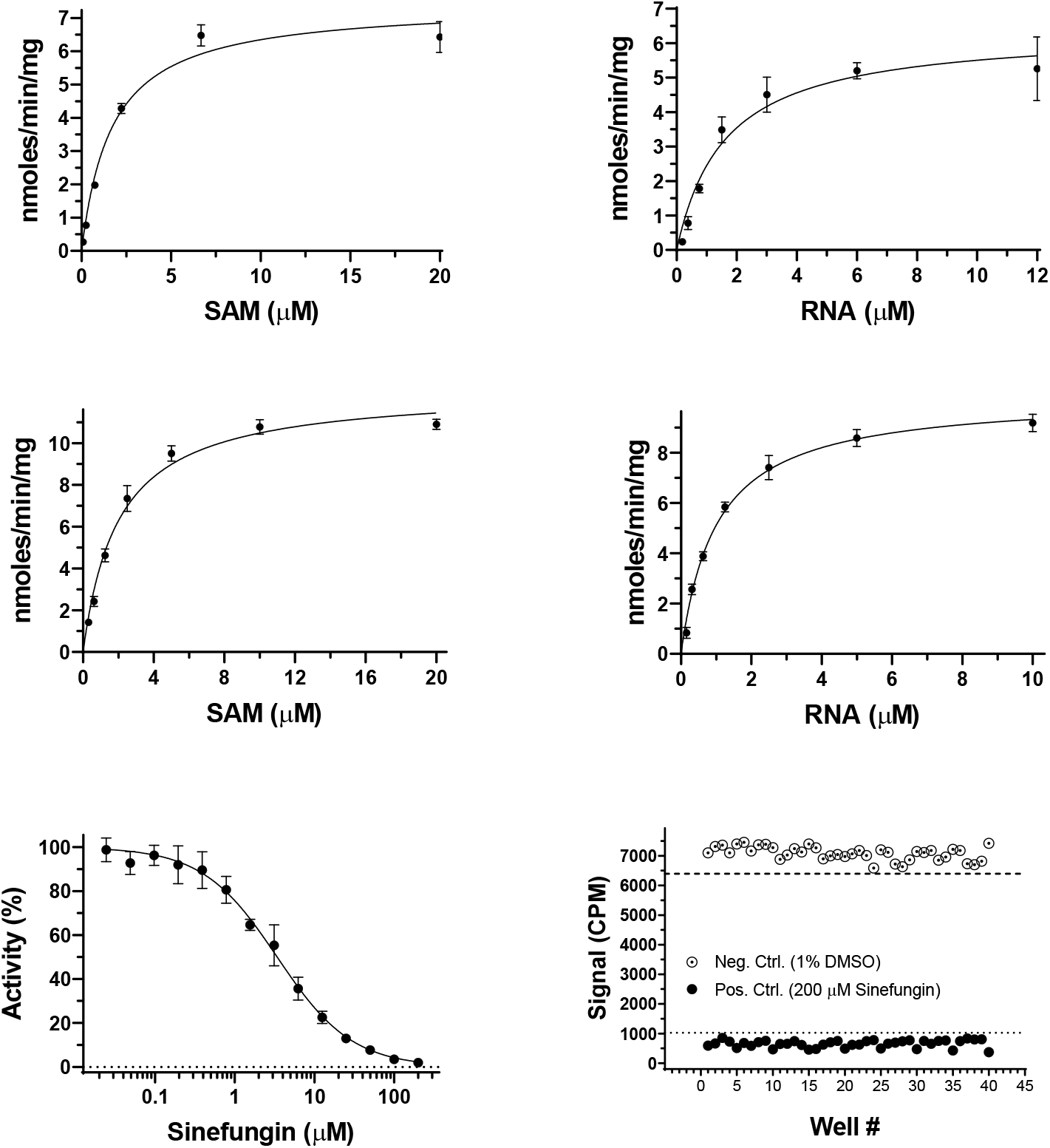
Kinetic parameter determination for nsp10-nsp16 complex. *K*_*m*_ values of nsp10-nsp16 complex were determined for SAM **(A** and **C)** and RNA substrate (**B** and **D**) under the optimized assay condition using a radiometric MTase assay as described in the experimental section at 250 nM (**A, B**) and 125 nM (**C, D**) of protein complex. (**E**) Sinefungin inhibited nsp10-nsp16 activity with an IC_50_ of 3.4 ± 0.4 μM (Hill Slope: −0.9). Experiments (**A-E**) were performed in triplicate. (**F**) The Z′-factor was determined in the presence (filled black circles) and absence (empty circles) of 200 μM Sinefungin in 0.8 μM RNA, 1.7 μM SAM, and 125 nM nsp16.

### 3.3. Assessment of HTS-Screening Amenability

To assess the quality of the developed assay for HTS-screening, first the linearity of the reaction over time under the screening conditions was analysed (**Suppl. Fig. 2D**). The time-course experiments revealed that the reaction was linear for at least 30 min. Using this assay condition, it was shown that sinefungin inhibited nsp10-nsp16 activity with an IC_50_ of 3.4 ± 0.4 μM (Hill Slope: −0.9) (**Fig. 2E**). Subsequently, the quality and robustness of the developed assay for high-throughput screening was analyzed. For screening in a 384-well format, a Z′-factor of 0.83 was attained (**Fig. 2F**). The optimized assay was then employed to screen a panel of 76 epigenetic chemical probes (**Suppl. Fig. 3)**, which included more than 20 MTase inhibitors **(Suppl. Table 1**). At a final compound concentration of 50 μM, none of these highly selective compounds significantly inhibited (>26%) the activity of nsp10-nsp16 complex, while SAH (IC_50_ of 5.9 ± 0.6 μM; **Suppl. Fig. 1F**), reduced the activity of nsp10-nsp16 by >90% at 50 μM.

**Figure 3.**
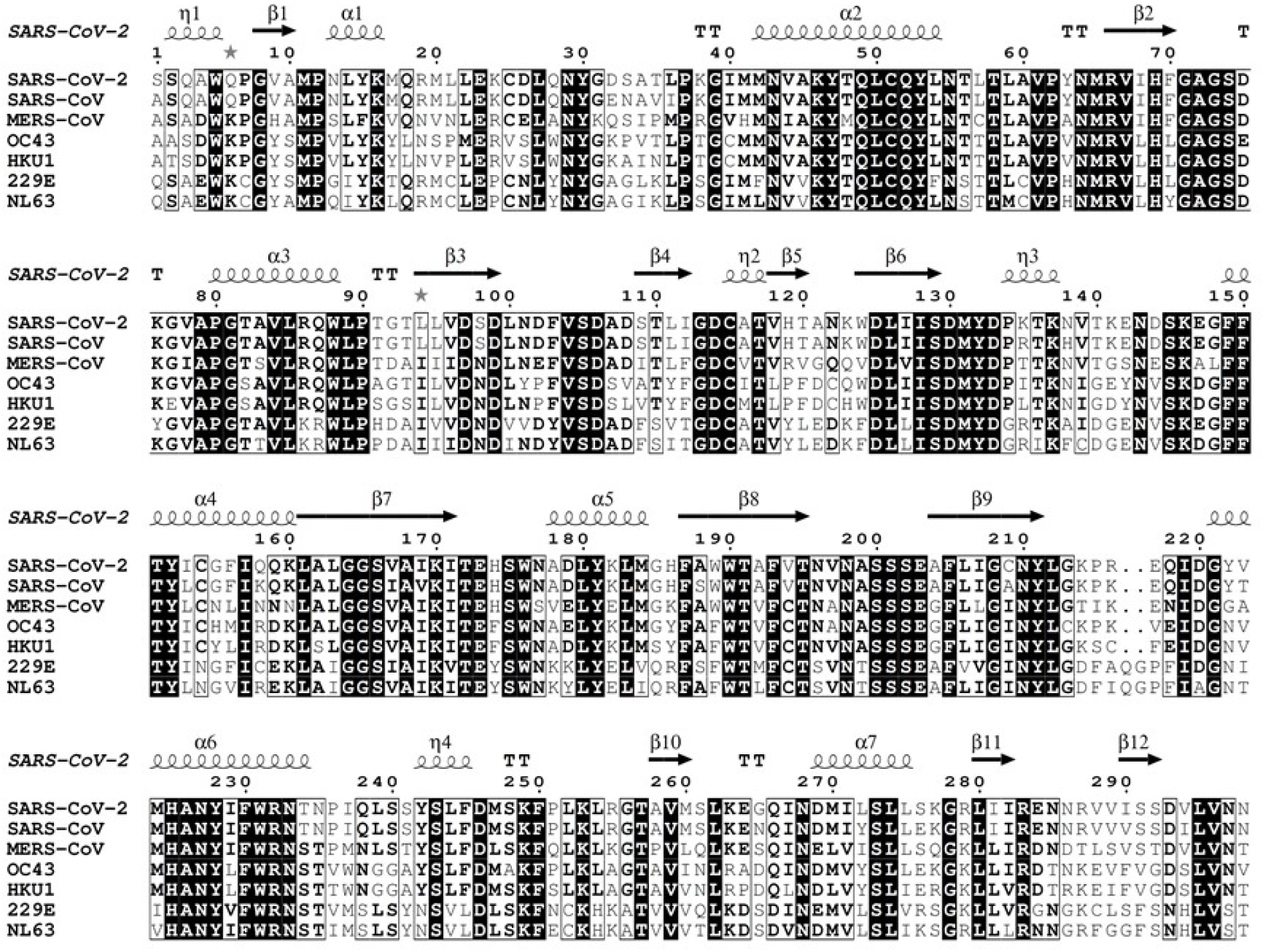
Sequence alignment of nsp16 from pathogenic CoVs. Amino acid sequences of nsp16 from 7 pathogenic CoVs (HKU1, NL63, OC43, 299E, MERS-CoV, SARS-CoV, and SAR-CoV-2) were aligned using Clustal Omega, and sequence similarities and secondary structure features were rendered by ESPript 3.0. Crystal structure of SAR-CoV-2 nsp10-nsp16 (PDB: 7JHE) was employed for extracting the secondary structure information. SARS-CoV-2 nsp16 shows 57.05, 58.39, 66.11, 63.76, 66.11, and 93.29% sequence identity to 299E, NL63, OC43, HKU1, MERS-CoV, SARS-CoV, respectively.

## 4. Discussion

As the fight against COVID-19 continues, several vaccines against SARS-CoV-2 have been made available to public. However, administering these vaccines requires very specific handling protocols, such as extremely low storage temperature for some, which may not be easily achievable in many countries. Even if all conditions are met, it will take many months to complete the vaccination. In addition, these vaccines may not be effective on fast mutating coronaviruses. This necessitates antiviral development ^36^. The 2′-O-MTase nsp16 has been proposed as an appealing target for development of anti-coronaviral therapeutics ^8, 11,28, 37^. Deletion of SARS-CoV nsp16 coding-region resulted in a blockade of viral RNA synthesis ^18^, and nsp16 mutants have shown a strong attenuation in infected mice ^19^. It has been suggested that nsp10-nsp16 complex, through its mRNA-capping activity, helps the CoVs evade the host immune system ^15^, therefore, any interruption in the activity of the nsp10-nsp16 could hinder the pathogenesis of CoVs through eliciting an immune response ^13,15, 16^. Inhibition of nsp10-nsp16 complex MTase activity by SAH (the product of the reaction), sinefungin (a SAM analogue) and aurintricarboxylic acid have been reported ^22, 23, 38^. However, potent and cell-permeable nsp10-nsp16 inhibitors are yet to be developed. The availability of activity-based HTS-screening assays would greatly enabe drug discovery. Activity of SARS-CoV nsp10-nsp16 complex has previously been assessed using a filter binding-based assay ^22^. Most recently, an HTS RNA-displacement assay has been reported for SARS-CoV-2 nsp10-nsp16 complex that will detect RNA competitive inhibitors.^31^ The nsp10-nsp16 complex activity assays reported to-date are low throughput ^22-24, 38^.

Here we reported development of a radioactivity-based assay for screening SARS-CoV-2 nsp10-nsp16 complex in a 384-well format. Since around 10-fold molar excess of nsp10 is required for the maximum *in vitro* MTase activity of nsp16 ^23^, a 1:8 ratio of nsp16 to nsp10 was chosen to ensure a near maximum activity of the complex. The kinetic parameters of nsp10-nsp16 complex methyltransferase activity are presented for the first time. Thus, the *K*_*m*_ of SAM and RNA were determined to be 2.0 ± 0.2 μM and 1.0 ± 0.1 μM, respectively. The ITC *K*_*d*_ values of 5.59 ± 1.15 μM and 1.21 ± 0.41 μM for SAM and RNA, respectively, were previously reported for nsp10-nsp16 complex from SARS-CoV ^24^. The IC_50_ values for SAH and sinefungin determined in this study were consistent with previously reported values for SARS-CoV and MERS-CoV nsp10-nsp16 complex.^22, 23^ Testing a subset of potent and selective chemical probes for human methyltransferases did not significantly inhibit the nsp10-nsp16 complex activity, indicating that the assay has a very low rate of false positives and is well suited for HTS. Unlike RNA displacement assays, this methyltransferase activity assay is suitable in detecting both SAM- and RNA competitive inhibitors.

The available evidence indicates that many other CoVs currently in various animals are preadapted to likely infect humans in some point of time in the future and cause new pandemics ^39-41^. Considering the natural diversity of CoVs across the globe ^1^ and the close interactions of humans with wild and domesticated animals, these future pandemics may not be prevented by the current vaccines ^36^. This further highlights the importance of developing potent inhibitors against coronaviral proteins that are conserved across this family of viruses toward developing pan-coronavirus therapeutics. Nsp16 is highly conserved across the CoV family ^12^, and available structures from several coronaviral species also reveal a high degree of structural conservation ^24, 26-28,30^. For example, SARS-CoV-2 nsp16 shows a minimum sequence identity of 57.05 % with the other pathogenic CoVs (**Fig. 3**). Mapping this sequence alignment on the nsp10-nsp16 structure (**Fig. 4**) demonstrates the conservation of SAM- and RNA-binding pockets across CoV species. Therefore, inhibitors targeting the active-site of nsp10-nsp16 may be effective against other emerging and re-emerging CoV strains. The radioactivity-based assay reported here will be an enabling tool towards developing such pan inhibitors of nsp10-nsp16 methyltransferase activities and possibly future pan-coronavirus therapeutics.

**Figure 4.**
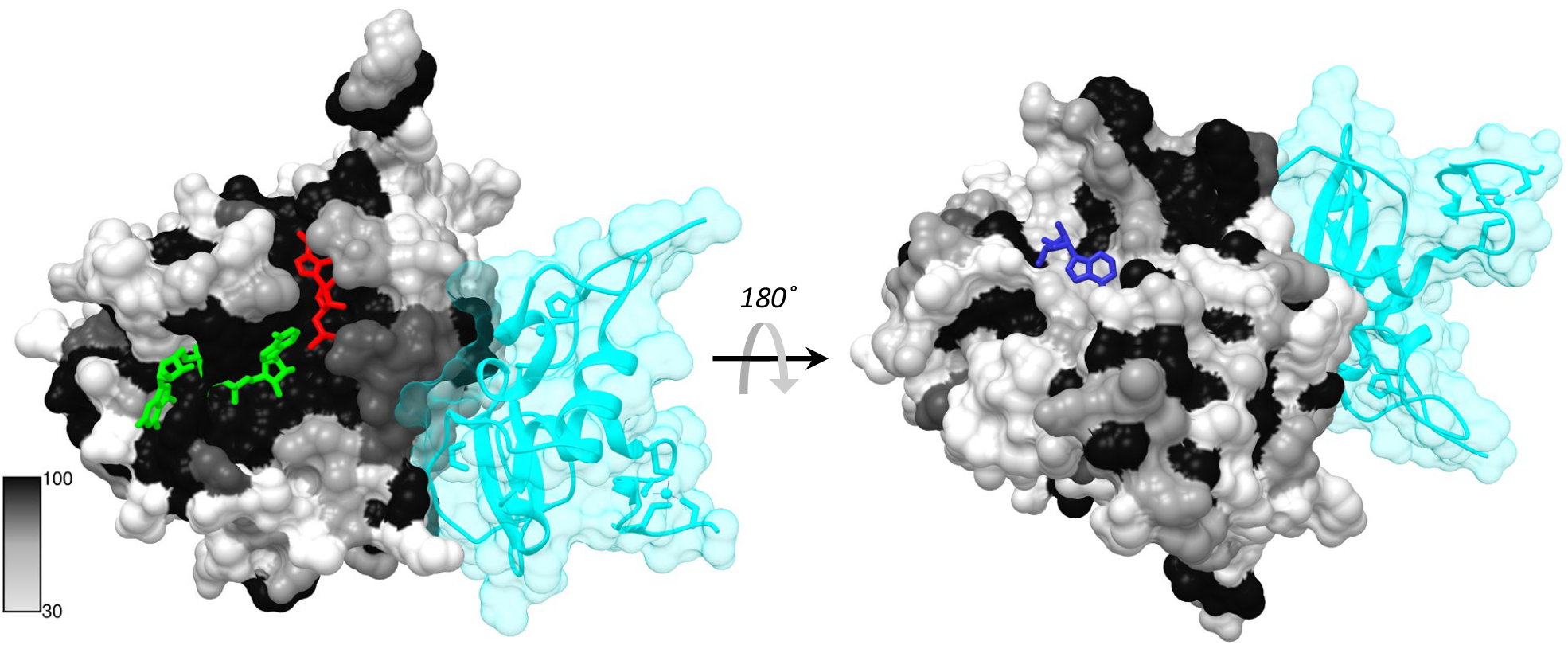
Sequence conservation of nsp16 across CoV species. Sequence conservation shown in Figure 3 is mapped onto the crystal structure of nsp10-nsp16 from SARS-CoV-2 (PDB: 6WKS). The structure was rendered by the percentage conservation of amino acid resides across the nsp16 from 7 human pathogenic CoVs (darker colors represents a higher degree of conservation, with black being the highest). Nsp10 subunit is shown in transparent cyan. SAM is shown with red stick, RNA is in green, and adenosine is represented by blue stick.

## 5. Conclusion

An HTS assay for assessing the activity of SARS-CoV-2 nsp10-nsp16 complex using a SPA-based method was developed. This assay provides a robust and sensitive tool for screening large libraries of compounds and is suitable for identifying inhibitors with different mechanisms of inhibition. It can be employed as an orthogonal method for re-evaluating potential inhibitors identified through other biochemical, biophysical, or cellular screening methods. Considering the critical role of nsp10-nsp16 complex in coronaviral pathogenesis and the highly conserved nature of nsp10-nsp16 complex across CoV species, the identified inhibitors may prove effective against other pathogenic CoVs, preventing future pandemics.

## Acknowledgement

We thank Dr. Aled Edwards and Dr. Cheryl Arrowsmith for continued support, and Dr. Peter Brown for critical review of the manuscript. This research was funded by the University of Toronto COVID-19 Action Initiative-2020, Takeda California, Inc., and COVID-19 Mitacs Accelerate postdoctoral awards to A.K.Y and S.P. The Structural Genomics Consortium is a registered charity (no: 1097737) that receives funds from; AbbVie, Bayer Pharma AG, Boehringer Ingelheim, Canada Foundation for Innovation, Eshelman Institute for Innovation, Genentech, Genome Canada through Ontario Genomics Institute [OGI-196], EU/EFPIA/OICR/McGill/KTH, Diamond Innovative Medicines Initiative 2 Joint Undertaking [EUbOPEN grant 875510], Janssen, Merck KGaA (aka EMD in Canada and US), Merck & Co (aka MSD outside Canada and US), Pfizer, São Paulo Research Foundation-FAPESP, Takeda and Wellcome [106169/ZZ14/Z].

## Author contributions

A.K.Y designed and performed experiments, analyzed data, and wrote the manuscript, F.L performed reproducibility confirmation experiments, F.L, K.D, and S.P contributed to assays, P.G and T.H purified proteins and prepared the protein complex, A.B contributed to compound management, M.V was the Principal Investigator, conceived the idea, lead the overall project, designed experiments, reviewed data, and wrote the manuscript. All authors reviewed the manuscript.

---

**Table1.** Kinetic characterization of nsp10-nsp16 complex.

| **Experiment** | **nsp16 (nM)** | **Substrate** | **$K_m^{app}$ (μM)** | **$k_{cat}$ ($\text{h}^{-1}$)** |
| --- | --- | --- | --- | --- |
| **1** | **250** | **SAM** | **$1.7 \pm 0.3$** | **$15.9 \pm 1.2$** |
|  |  | **RNA** | **$1.6 \pm 0.4$** |  |
| **2** | **125** | **SAM** | **$2.0 \pm 0.2$** | **$26.9 \pm 0.3$** |
|  |  | **RNA** | **$1.0 \pm 0.1$** |  |

## Supplementary Data

**Supplementary Figure 1.**
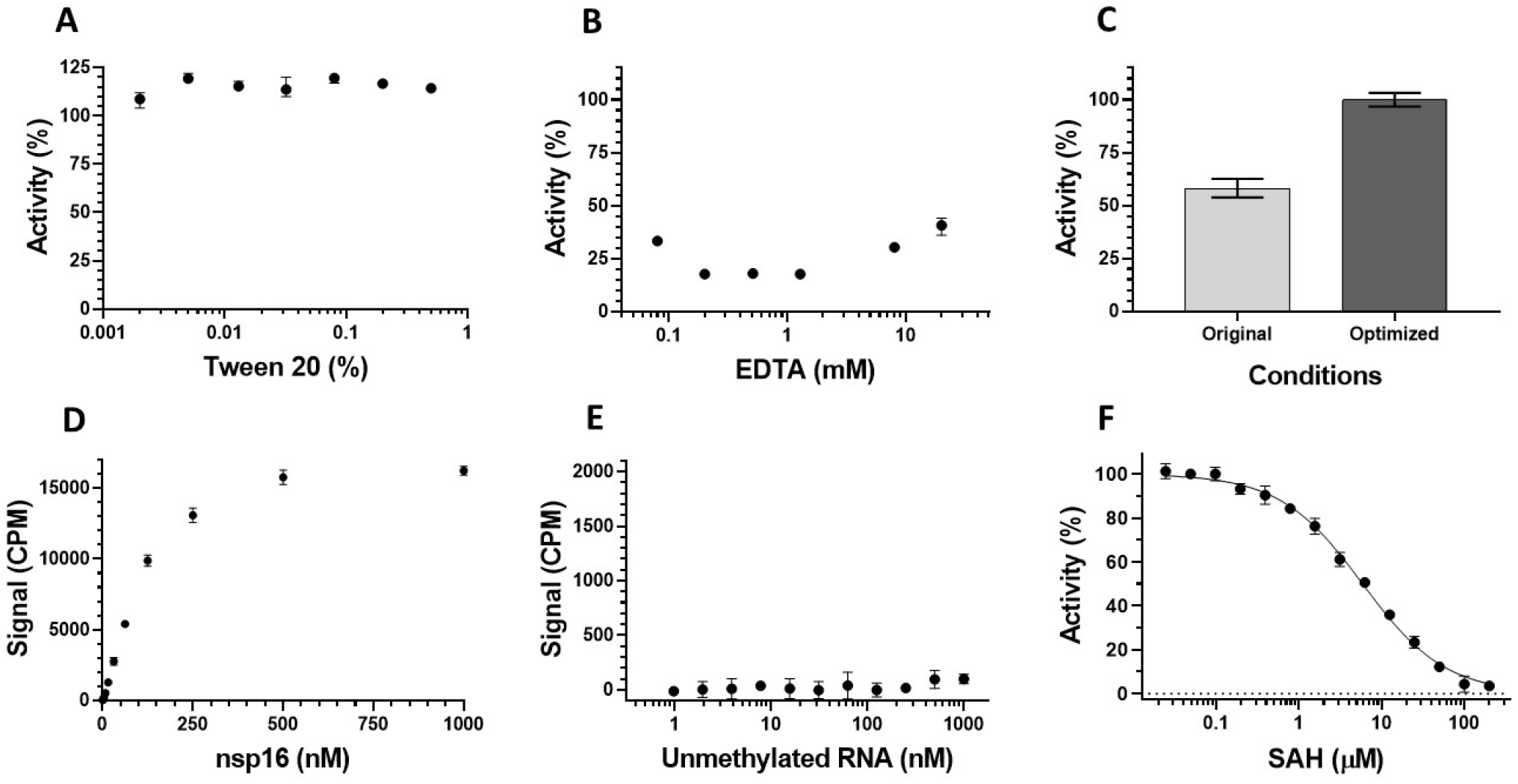
**(A)** Effect of Tween-20 as an additive on the activity of nsp10-nsp16 complex. **(B)** The inhibitory effect of EDTA on nsp10-nsp16 at various concentrations. **(C)** Comparison of nsp10-nsp16 MTase activity in the original buffer (50 mM Tris pH 8.0, 1 mM MgCl_2_, and 5 mM DTT) versus the optimized buffer condition (50 mM Tris pH 7.5, 100 mM KCl, 1.5 mM MgCl_2_, 5 mM DTT, 0.01% BSA, 0.01% Triton X-100) in the presence of 2 μM RNA substrate, 5 μM SAM (16% ^3^H-SAM), and 250 nM nsp16. **(D)** MTase activity at various concentrations of nsp10-nsp16 complex using N7-meGpppACCCCC RNA (Cap-0). **(E)** The N7-unmethylated RNA is not a substrate for nsp10-nsp16 complex; here reactions were performed in the presence of 125 nM nsp10-nsp16, 5 µM SAM and varying concentrations of N7-unmethylated RNA substrate (0.97 nM to 1 µM) for 30 minutes. **(F)** SAH inhibited nsp10-nsp16 with an IC_50_ value of 5.9 ± 0.6 μM (Hill Slope: −0.9); Experiments were performed in triplicate (n=3).

**Supplementary Figure 2.**
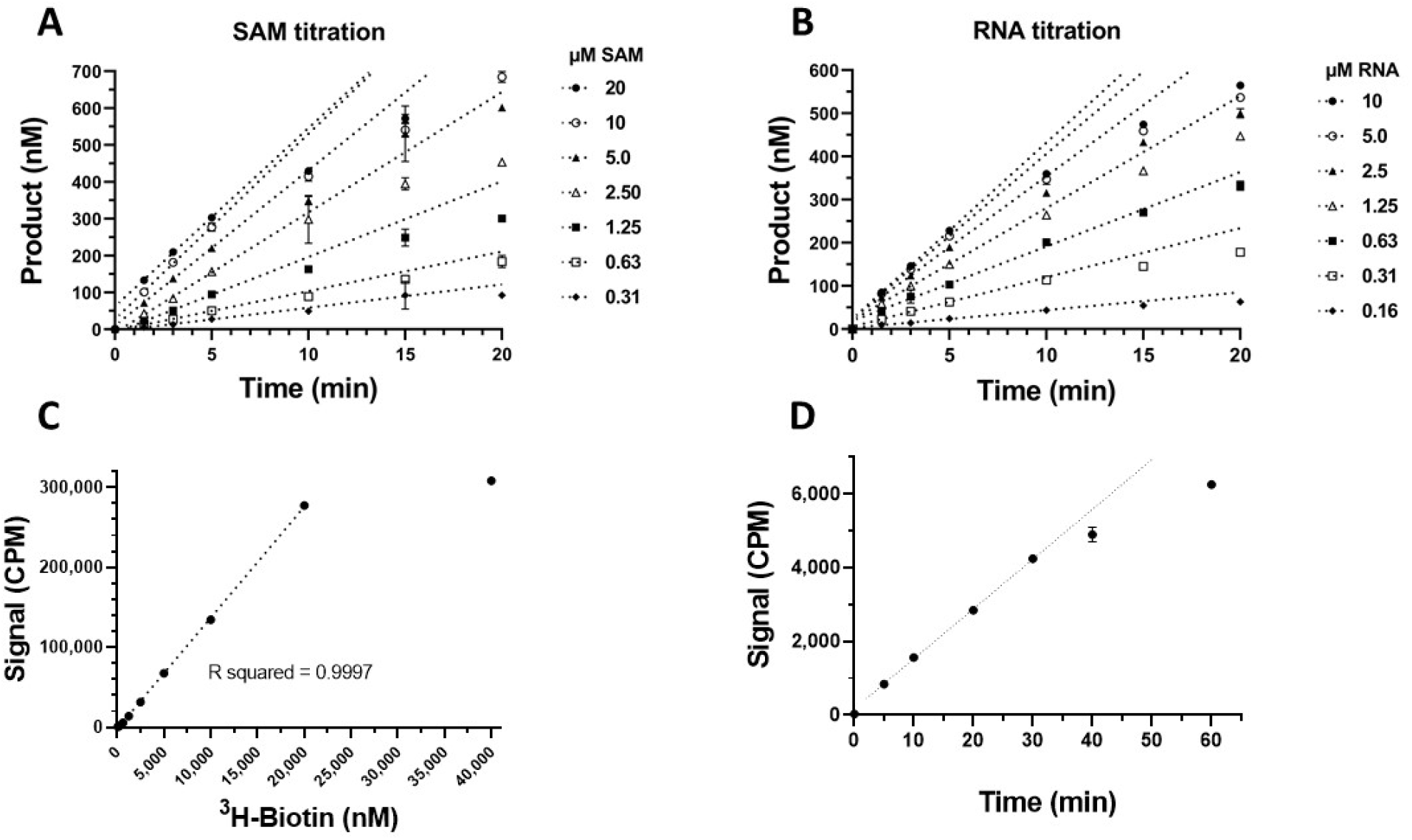
Time-based experiments for measuring the rates of reaction in the presence of 125 nM nsp16 for **(A)** SAM-titration experiments and **(B)** RNA-titrations. For all the experiments in A and B, 3 μL of reaction was added per well of the 96-well FlashPlate, and only the first 20 minutes of the reaction progress curve is shown here for clarity. **(C)** Titration of ^3^H-Biotin. 3 μL of ^3^H-Biotin was added per well of the 96-well SPA plate. **(D)** Testing the linearity of nsp10-nsp16 activity over time using the optimized buffer and screening conditions (i.e., 10 μL reactions containing 0.8 μM RNA, 1.7 μM SAM (33% ^3^H-SAM), and 125 nM nsp16 enzyme).

**Supplementary Figure 3.**
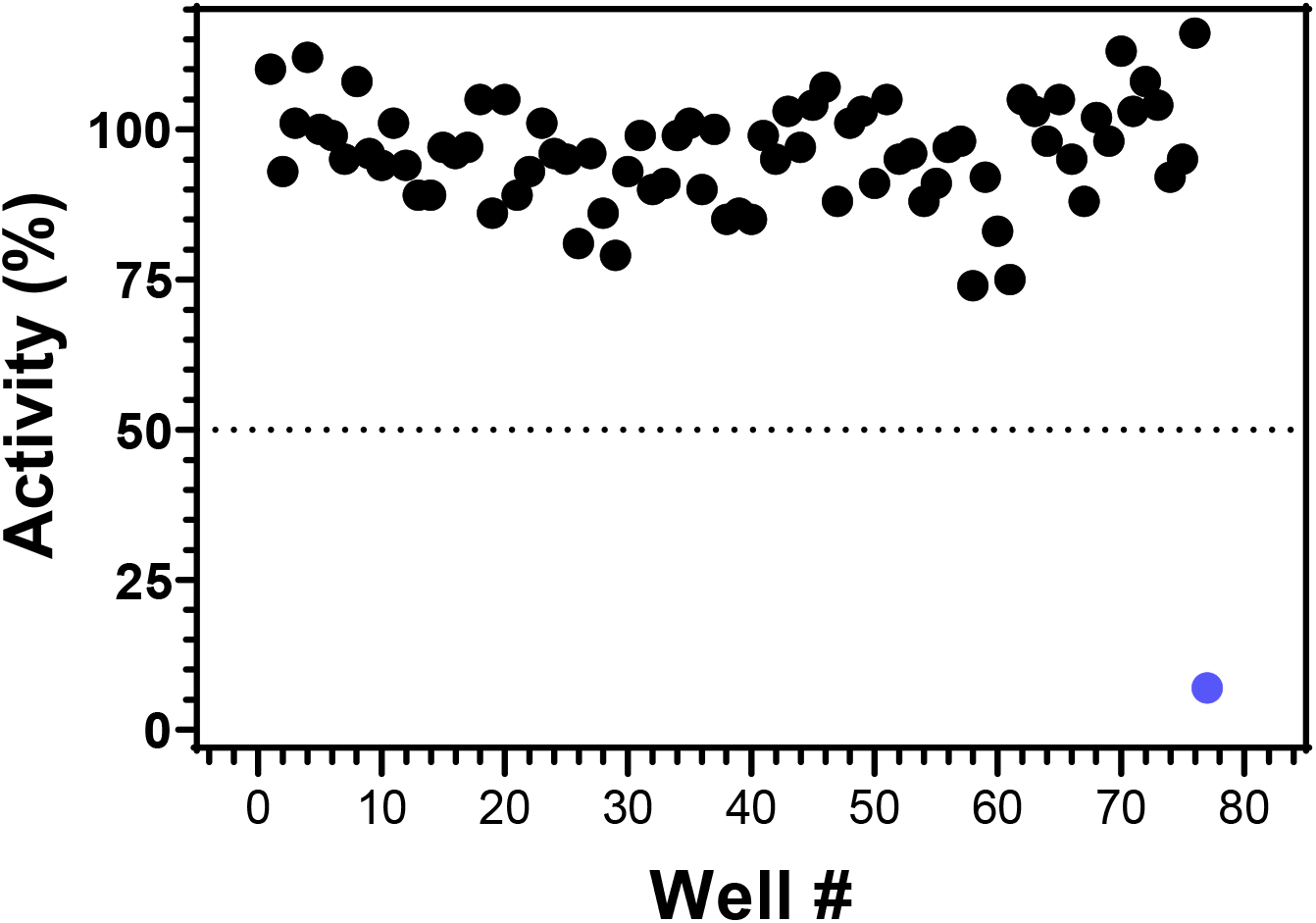
**Screening of 76 epigenetics probes and their analogous against nsp10-nsp16 complex.** A collection of 76 small-molecules were screened at 50 μM against nsp10-nsp16 under the developed screening assay conditions (i.e., 0.8 μM RNA, 1.7 μM SAM, and 125 nM nsp16 enzyme). The corresponding percentage activity data for each probe is shown on the graph with a black dot. SAH was used at a similar concentration as a control (blue dot). Please note that the dotted line marks the 50% activity threshold.

**Supplementary Table 1.** 76 epigenetic compounds were screened against nsp10-nsp16 complex. 76 compounds, including epigenetic probes and their closely related analogues, were screened against SARS-CoV-2 nsp10-nsp16 using the developed HTS assay. The observed percentage of activity of nsp10-nsp16 in the presence of each of these compounds (at 50 µM) is presented. The list of compounds (available at https://www.thesgc.org/chemical-probes), and their specific protein targets is provided. Negative control analogues of the chemical probes are specified with “Negative Ctrl” under the “Specific Targets” column.

| **Compound** | **Specific Targets** | **% activity**<br>**@ 50 μM** |
| --- | --- | --- |
| **UNC1215** | L3MBTL3 | 103 |
| **BSP** | pan-Bromodomain | 105 |
| **UNC0638** | EHMT2 (G9a), EHTM1 (GLP) | 102 |
| **UNC0737** | EHMT2 (G9a), EHTM1 (GLP)<br>(Negative Ctrl) | 113 |
| **A-395** | EED | 100 |
| **A-395N** | EED (Negative Ctrl) | 99 |
| **TP-064** | CARM1 (PRMT4) | 98 |
| **TP-064N** | CARM1 (PRMT4) (Negative<br>Ctrl) | 105 |
| **MS023** | Type I PRMTs (PRMT1,3,4,6,8) | 99 |
| **MS094** | Type I PRMTs (PRMT1,3,4,6,8)<br>(Negative Ctrl) | 103 |
| **UNC1999** | EZH2 | 108 |
| **UNC2400** | EZH2 (Negative Ctrl) | 104 |
| **PFI-2** | SETD7 | 105 |
| **(S)-PFI-2** | SETD7 (Negative Ctrl) | 110 |
| **SGC-CBP30** | CREBBP, EP300 | 103 |
| **A-366** | EHMT2 (G9a), EHMT1 (GLP) | 112 |
| **OICR-9429** | WDR5 | 103 |
| **OICR-0547** | WDR5 (Negative Ctrl) | 101 |
| **NVS-PAK1-1** | PAK1 | 107 |
| **GSK864** | Mutant isocitrate dehydrogenase<br>1 | 81 |
| **BI-9321** | NSD3 | 97 |
| **BI-9466** | NSD3 (Negative Ctrl) | 105 |
| **SGC707** | PRMT3 | 105 |
| **XY1** | PRMT3 (Negative Ctrl) | 116 |
| **PFI-3** | SMARCA2, SMARCA4,<br>PBRM1 (PB1) | 95 |
| **GSK-J1** | KDM6B (JMJD3), KDM6A<br>(UTX), KDM5B (JARID1B) | 96 |
| **A-485** | p300, CBP | 95 |
| **A-486** | p300, CBP (Negative Ctrl) | 108 |
| **SGC0946** | DOT1L | 97 |
| **SGC0649** | DOT1L (Negative Ctrl) | 91 |
| **UNC0642** | EHMT2 (G9a), EHTM1 (GLP) | 98 |
| **GSK343** | EZH2 | 93 |
| **L-Moses** | KAT2B (PCAF), KAT2A<br>(GCN5) | 100 |
| **GSK484** | PAD-4 | 101 |
| **BAY-876** | GLUT1 | 97 |
| **BAY-588** | GLUT1 (Negative Ctrl) | 94 |
| **PFI-5** | SMYD2 | 88 |
| **NVS-CECR2-1** | CECR2 | 104 |
| **OF-1** | BRPF1, BRD1 (BRPF2), BRPF3 | 88 |
| **IOX1** | pan-2-OG | 99 |
| **I-BRD9** | BRD9 | 79 |
| **LP99** | BRD9, BRD7 | 85 |
| **NI-57** | BRPF1, BRD1<br>(BRPF2), BRPF3 | 97 |
| **I-CBP112** | CREBBP, EP300 | 93 |
| **GSK-LSD1** | KDM1A (LSD1) | 86 |
| **GSK2801** | BAZ2A, BAZ2B | 89 |
| **BAZ2-ICR** | BAZ2A, BAZ2B | 96 |
| **PFI-4** | BRPF1B | 96 |
| **A-196** | SUV420H1/H2 | 93 |
| **A-197,**<br>**SGC2043** | SUV420H1/H2<br>(Negative Ctrl) | 101 |
| **GSK591** | PRMT5 | 96 |
| **SGC2096** | PRMT5 (Negative Ctrl) | 98 |
| **IOX2** | pan-2-OG | 90 |
| **PFI-1** | BRD2, BRD3, BRD4,<br>BRDT (BET) | 91 |
| **JQ1** | BRD2, BRD3, BRD4,<br>BRDT (BET) | 91 |
| **TP-472** | BRD9, BRD7 | 88 |
| **MS049** | PRMT4 (CARM1),<br>PRMT6 | 95 |
| **BAY-299** | BRD1, TAF1 | 96 |
| **TP-238** | CECR2, BPTF (FALZ) | 95 |
| **LLY-507** | SMYD2 | 90 |
| **BAY-598** | SMYD2 | 89 |
| **BAY-369** | SMYD2 (Negative Ctrl) | 94 |
| **BI-9564** | BRD9, BRD7 | 86 |
| **GSK6853** | BRPF1 | 95 |
| **SGC6870** | PRMT6 | 83 |
| **SGC6870N** | PRMT6 (Negative Ctrl) | 75 |
| **MRK-740** | PRDM9 | 86 |
| **MRK-740-NC** | PRDM9 (Negative Ctrl) | 85 |
| **SGC3027** | PRMT7 | 74 |
| **SGC3027N** | PRMT7 (Negative Ctrl) | 92 |
| **UNC6934** | NSD2-PWWP1 | 92 |
| **UNC7145** | NSD2-PWWP1<br>(Negative Ctrl) | 95 |
| **LLY-283** | PRMT5 | 99 |
| **LLY-284** | PRMT5 (Negative Ctrl) | 101 |
| **BAY-6035** | SMYD3 | 89 |
| **BAY-444** | SMYD3 (Negative Ctrl) | 101 |
| **SAH**<br>**(Control)** | - | 7 |

## Notes

### Competing Interest Statement

The authors have declared no competing interest.

